# Protein arginine-methyltransferase 1 (PRMT1): a new pharmacological target in cholangiocarcinoma

**DOI:** 10.64898/2026.05.29.728163

**Authors:** Emiliana Valbuena-Goiricelaya, Jasmin Elurbide, Maria Ujue Latasa, Amaya Lopez-Pascual, Iker Uriarte, Leticia Colyn, Patricia Inacio, Robert Arnes-Benito, Elena Adan-Villaescusa, Borja Castello-Uribe, Barbara Franceschini, Flavio Milana, Pavel Strnad, Sona Frankova, Eva Sticová, Ondrej Fabian, Irene Amat, Jesus Urman, Ana Lleo, Meritxell Huch, Maria Arechederra, Carmen Berasain, Maite G Fernandez-Barrena, Matias A Avila

**Affiliations:** Hepatology Laboratory, Solid Tumors Program, CIMA, CCUN, University of Navarra, Pamplona, Spain; CIBERehd, Madrid, Spain; Instituto de Investigaciones Sanitarias de Navarra IdiSNA, Pamplona, Spain; Max Plank Institute of Molecular Cell Biology and Genetics, Dresden, Germany; Hepatobiliary Immunopathology Lab, IRCCS Humanitas Research Hospital, Rozzano, Italy; Division of Hepatobiliary and General Surgery, IRCCS Humanitas Research Hospital, Rozzano, Italy; Department of Biomedical Sciences, Humanitas University, Pieve Emanuele, Italy; Department of Gastroenterology and Hepatology, Medical University Lausitz - Carl Thiem (MUL-CT), Cottbus, Germany; Department of Hepatogastroenterology, Institute for Clinical and Experimental Medicine, Prague, Czechia; Clinical and Transplant Pathology Centre, Institute for Clinical and Experimental Medicine, Prague, Czechia; Department of Pathology, 3rd Faculty of Medicine, Charles University, and Kralovske Vinohrady University Hospital, Prague, Czechia; Department of Pathology, Navarra University Hospital, Pamplona, Spain; Department of Digestive Diseases, Navarra University Hospital, Pamplona, Spain; Division of Internal Medicine and Hepatology, Department of Gastroenterology, IRCCS Humanitas Research Hospital, Rozzano, Italy

**Author notes:** Corresponding author. Dr. Matias A Avila, Hepatology Laboratory, Solid Tumors Program, CIMA, CCUN, University of Navarra, Pamplona, Spain. These authors share first authorship (Emiliana Valbuena-Goiricelaya, Jasmin Elurbide, Maria Ujue Latasa). These authors are co-senior authors of this study (Maite G Fernandez-Barrena, Matias A Avila).

**Keywords:** Cholangiocarcinoma, arginine methylation, epigenetic therapy, immunotherapy

## Abstract

Cholangiocarcinoma (CCA) is a highly aggressive malignancy characterized by poor prognosis, limited therapeutic options, and a predominantly immunosuppressive tumor microenvironment. Protein arginine methyltransferase 1 (PRMT1), the major mediator of asymmetric arginine dimethylation, has been implicated in multiple oncogenic processes, although its role in CCA remains unknown. Here, we demonstrate that PRMT1 is frequently overexpressed in human CCA and is associated with aggressive molecular subtypes and immune-desert tumors. Genetic dependency analyses and pharmacological inhibition using type I PRMT inhibitors markedly impaired CCA cell proliferation, clonogenicity, and tumoroid growth. Transcriptomic profiling revealed that PRMT1 inhibition induces broad alterations in gene expression and alternative splicing, affecting pathways involved in proliferation, apoptosis, DNA damage response, metabolism, and immune signaling. Mechanistically, PRMT1 targeting promoted genomic stress, accumulation of cytosolic double-stranded DNA, and activation of the cGAS-STING-TBK1-IRF3 signaling axis, resulting in enhanced interferon signaling and increased expression of T cell-recruiting chemokines, including CXCL9 and CXCL10. PRMT1 inhibition also synergized with cisplatin, poly-ADP-ribose polymerase (PARP) inhibition, and PRMT5 blockade in vitro and in patient-derived tumoroids. Importantly, in an aggressive orthotopic murine model of intrahepatic CCA, combined treatment with the PRMT1 inhibitor GSK3368715 and anti-PD-1 antibodies significantly reduced tumor burden and increased CD4+ and CD8+ T-cell infiltration compared with monotherapies. Collectively, these findings identify PRMT1 as a critical regulator of CCA growth and immune evasion and support the therapeutic potential of PRMT1 inhibition, particularly in combination with immunotherapy.

## 1. Introduction

Cholangiocarcinoma (CCA) is the second most common primary liver malignancy[1]. CCAs arise from distinct anatomical locations, leading to intrahepatic (iCCA) and extrahepatic (eCCA) subtypes, this latter with perihilar pCCA and distal dCCA variants, each exhibiting unique histological and molecular features[2–5]. Although geographical variations exist, the global incidence and mortality of CCA, particularly iCCA, have risen in recent years. Notably, the steepest increase has occurred among younger individuals (18-44 years), and mortality rates are two- to three-fold higher in men than in women across most regions[6,7]. The majority of CCAs are diagnosed at advanced stages, when curative interventions such as surgical resection or liver transplantation are no longer feasible. The combination of late diagnosis and the aggressive biological behavior of these tumors results in a dismal 5-year overall survival rate of less than 10%[1]. For advanced disease, CCA exhibits limited responsiveness to conventional chemotherapy regimens, such as cisplatin plus gemcitabine [1]. Targeted agents against actionable alterations, such as *IDH1* mutations or *FGFR2* fusions and rearrangements, have shown promising results in genotype-selected patients, although these are a minority of cases, and acquired resistance may develop[8–10]. With the exception of a small subset of patients harboring microsatellite instability-high tumors, responses to immune checkpoint inhibitors (ICIs) as monotherapy have been disappointing[7]. However, recent clinical trials have demonstrated that combining conventional chemotherapy with ICIs improves therapeutic efficacy, establishing chemoimmunotherapy as the current first-line standard of care[7,8,11]. Despite these advances, the overall prognosis for patients with CCA remains poor[7,12]. Further exploration of novel therapeutic combinations is warranted to overcome the predominantly immunosuppressive tumor microenvironment (TME) in CCA and enhance patient outcomes[8,11,13–15].

Epigenetic dysregulation plays a key role in CCA initiation and progression, encompassing widespread DNA methylation alterations and histone post-translational modifications (PTMs) beyond chromatin-remodeling gene mutations [16]. Because epigenetic alterations are reversible, epigenetic therapies represent promising strategies for cancer treatment [17,18], including CCA, as supported by the antitumor activity of small-molecule inhibitors targeting DNA- and histone lysine-methyltransferases in experimental CCA models. [19–22]. Importantly, emerging precinical evidence indicates that specific epigenetic changes may modulate tumor-TME interactions influencing antitumor immune responses, and that their pharmacological inhibition can enhance ICI’s efficacy [23–26].

Among epigenetic PTMs, arginine methylation represents an additional regulatory mechanism mediated by a family of nine protein arginine methyltransferases (PRMTs). These enzymes are categorized into three classes: Type I PRMTs (PRMT1, 2, 3, 4, 6, and 8), which catalyze asymmetric arginine dimethylation; Type II PRMTs (PRMT5 and PRMT9), responsible for symmetric dimethylation; and the Type III enzyme PRMT7, which generates monomethylated arginine residues [27]. PRMT dysregulation, predominantly driven by overexpression, is increasingly recognized as a contributor to cancer pathogenesis through widespread effects on signalling pathways, transcriptional programs, RNA metabolism, DNA damage responses, and tumor immunity [28–31]. In this context, we recently identified PRMT5 upregulation as a pro-tumorigenic event in CCA and demonstrated the antitumoral efficacy of its pharmacological inhibition in experimental models [32]. Besides PRMT5, overexpression of PRMT1 has been strongly linked to the development of a variety of cancer types. PRMT1 is responsible for 85% of asymmetric dimethylarginine (ADMA) marks on cellular proteins through the sequential methylation of one of the terminal guanidine nitrogen atoms. All the oncogenic mechanisms described above for PRMTs have also been demonstrated for PRMT1, with promotion of cell growth through modulation of gene expression, DNA damage repair, and evasion of antitumor immunity playing pivotal roles [33–36]. This critical involvement of PRMT1 in cancer development has spurred the development of inhibitors, such as GSK3368715 and MS023 that have shown potent antitumoral effects as single agents in mouse models of different tumor types, and promising results in combinatorial strategies with DNA damaging agents, ICIs and other epigenetic drugs [34,37–39]. In this study we show that PRMT1 is frequently overexpressed in CCAs, and we provide experimental *in vitro* and *in vivo* evidence supporting the efficacy of PRMT1 inhibition for the treatment of this malignancy. Our findings identify PRMT1 as a potential target to overcome imunotherapy resistance in CCA.

## 2. Methods

### 2.1. Human tissue samples

CCA tissues were obtained from patients with iCCA (n=86; 83 surgical samples, 3 needle-biopsies) or eCCA (n=23) that underwent biopsies or surgical resection for diagnostic or therapeutic purposes. Sample collection and use were approved by the institutionś human research committees: Institute for Clinical and Experimental Medicine and Thomayer University Hospital, Prague, Czech Republic (protocol # 40242/25, A-25-21), at the Hospital Universitario de Navarra Pamplona, Spain (ethical committee approval # 2015/120) and at Humanitas Research Hospital, Milan, Italy (Institutional Review Board approval # 146/20 and 776/22). Tumor grade was established upon anatomopathological examination of tissue sections according to the WHO Classification of Tumours, Digestive System Tumours, 5th edition.

### 2.2. Cell culture, treatments and reagents

The characteristics and origin of the CCA cell lines used in this study have been previously described [40]. The human cell lines HuCCT-1 and TFK-1 were cultured in Dulbecco’s Modified Eagle Medium (DMEM)-F12, while RBE cells were cultured in Roswell Park Memorial Institute (RPMI)-1640 medium, each containing 10% heat-inactivated Fetal Bovine Serum (FBS) and antibiotics (Gibco Thermo-Fisher, Waltham, MA, USA), following previously reported conditions [19]. Treatment times and dosages in the experiments are specified throughout the manuscript, with controls receiving equivalent concentrations of dimethyl sulfoxide (DMSO) (always < 0.1% of the final volume). When indicated, cells were stimulated with human IFNγ (R&D Systems, Minneapolis, MN, USA) for 24h and transfected with vaccinia virus dsDNA V70mer [41] using lipofectamine 2000 reagent (11668019, invitrogen).

For proliferation studies, cells were seeded at a density of 2500-3000 cells/well in 96-well plates (triplicates). After overnight incubation, cells were treated for seven days with vehicle alone (0.1% DMSO), and/or PRMT type I inhibitors GSK3368715, MS023 or PRMT1 specific inhibitor TCE-5003. Drugs were from MedChemExpress (Monmouth Junction, NJ, USA) and were added to cultures in complete medium. Experiments were repeated three times for each cell line. Cell viability was measured using the CellTiter 96 Aqueous One Solution Cell Proliferation Assay (Promega, Madison, WI, USA). All cells were routinely tested for Mycoplasma. The drug concentration required to inhibit cell growth by 50% compared to the untreated control (GI_50_) was calculated after curve fitting with GraphPad-Prism-v9 software as previously described [19]. For the calculation of combination index (CI) values, growth inhibition was determined at different combined concentrations of the PRMT1i and the antitumoral drugs cisplatin (cis-diamminedichloroplatinum II, CDDP) (Sigma Aldrich, St. Louis, MO, USA), Olaparib, the PRMT5 inhibitor GSK3326595 (GSK595), or the MTAP inhibitor methylthio-DADMe-immucillin-A (MTDIA) (MedChemExpress). Briefly, 3000 cells were cultured in 96-well plates in triplicates and the different concentrations of compounds and their combinations were added in a final volume of 100 µL. After 72h, viability was measured using the Cell Titer 96 Aqueous One Solution Cell Proliferation Assay. Data were analyzed using the Calcusyn software (Biosoft, Cambridge, UK), and the CI determined whether the effects of drug combinations were additive (CI=1) or synergistic (CI<1) as described [19].

For colony formation assays, 1,000 cells per well were seeded in 6-well plates. After 24h, GSK3368715 or an equivalent volume of DMSO (vehicle control) was added. At the end of the treatment period (7–10 days), plates were washed once with PBS, fixed with 4% formaldehyde, and stained with 0.5% crystal violet. Plates were photographed, and the dye was subsequently solubilized in 10% acetic acid. For quantification, absorbance was measured at 600 nm using a spectrophotometer, as previously described [20].

### 2.3. In vivo experiments

CCA was induced in 5 weeks-old C57BL/6J male mice by hydrodynamic tail vein injection (HTVI) of plasmids coding for mutant TAZ (TAZS89A, 10 μg/mouse), myr-AKT (10 mg/mouse) and the sleeping beauty transposase (SB, 0.8 mg/mouse) (GenScript, Piscataway, NJ, USA) as described [42]. One group of mice was treated with GSK3368715 (MedChemTronica) (75 mg/kg *p.o.* daily) another group was treated with anti-PD1 (RMP1-14, Bio X Cell, 100 μg per mouse) and another with the combination (n=6 per group). Anti-PD1 was administered via intraperitoneal injection once per week for three administrations. Control mice (n=6) received the same volume of vehicle, as described [26]. As reported for other HTVI oncogene-induced CCA models, treatments started two days after HTVI [21], and mice were treated for two or three weeks. C57BL/6J mice (n=4) without HTVI injection nor treatment were used as healthy controls and were sacrificed at the same ages as the treated groups. All animals received humane care and protocols were performed according to the guidelines of the Animal Care Committee of the University of Navarra (approval #R-CP001-15GN).

### 2.4. Immunohistochemical analyses

Immunostainings of human CCA tissue samples were performed as we previously described [32] using PRMT1 antibody from Abcam (Cambridge, UK, ab190892, 1:500 dilution). PRMT1 scores were calculated using QuPath software (v0.7.0) as reported [19]. For each whole-slide image, representative regions of interest (ROIs) were manually annotated in histologically confirmed tumour areas. Within each ROI, all tumor cell nuclei were segmented using QuPath’s Positive cell detection module based on the haematoxylin counterstain. A semi-quantitative composite score, conceptually analogous to a modified H-score was then derived as the product of two ordinal sub-scores. The proportion of positive tumour nuclei was stratified into four categories scored 0–3 (0: 0%; 1: 1–32%; 2: 33–66%; 3: >67%), and the mean nuclear DAB optical density was likewise stratified into four categories scored 0–3, corresponding to absent, weak, moderate, and strong staining. The final PRMT1 score per slide was computed as the product of the percentage sub-score and the intensity sub-score, yielding a composite index ranging from 0 to 9. Scoring was performed by a single observer blinded to clinicopathological data.

For *in vivo* studies immunohistochemistry was performed on 3-μm formalin-fixed, paraffin-embedded mouse liver tissue sections as previously described [32], using the following primary antibodies: anti-CD8 (98941, Cell Signaling Technology, Danvers, MA, USA, 1:800), anti-CD4 (ab183685, Abcam, 1:1000), anti-γH2AX (9718, Cell Signaling Technology, 1:1000), anti-STING (4947, Cell Signaling Technology, 1:1000), anti-phospho-IRF3 (13627, Cell Signaling Technology, 1:800). Briefly, antigen retrieval was performed in 10 mM Tris-EDTA buffer (pH 9), and detection was carried out with HRP-conjugated EnVision polymer (K400311-2, Dako) followed by DAB (K3468, Dako). Negative controls were run in parallel with omission of the primary antibody. Stained slides were digitised and subsequently analysed using QuPath software (v0.7.0).

### 2.5. Western Blot

Cells and tissues were lysed in RIPA buffer as described previously [43]. Homogenates form cells and tissues, were subjected to immunoblot (Western blot) analysis as reported [44]. Total protein was separated via sodium dodecyl sulfate polyacrylamide gel electrophoresis (SDS-PAGE), and electrotransferred onto PVDF membranes (Millipore, Burlington, MA, USA). The following primary antibodies were used for protein detection: anti-Symmetric Di-Methyl Arginine Motif (SDMA) (13222, Cell Signaling Technology, 1:1000), anti-Asymmetric Di-Methyl Arginine Motif (ADMA) (13522, Cell Signaling Technology, 1:1000), anti-Mono-Methyl Arginine (MMA) (8015, Cell Signaling Technology, 1:1000), anti-methylthioadenosine phosphorylase (MTAP) (4158S, Cell Signaling Technology, 1:1000), anti-TBK1 (3013, Cell Signaling Technology, 1:1000), anti-phospho-TBK1 (5483, Cell Signaling Technology, 1:1000), anti-IRF3 (4302, Cell Signaling Technology, 1:1000), pIRF3 (4947, Cell Signaling Technology, 1:1000), anti-STING (13647, Cell Signaling Technology, 1:1000), p-STING (PA-105674, Invitrogen, 1:1000), anti-DNA Methyltransferase 1 (DNMT1) (5032S, Cell Signaling Technology. 1:1000). Blots were probed with anti-α-Tubulin (2144, Cell Signaling Technology. 1:1000), anti-Heat Shock Protein 90 (HSP90) (4874S, Cell Signaling Technology: 1:1000), or anti-glyceraldehyde-3-phosphate dehydrogenase (GAPDH) (2118S, Cell Signaling Technology: 1:1000) to demonstrate equal loading. Secondary HRP-conjugated goat anti-mouse IgG (68860, Cell Signaling Technology, 1:5000) or anti-rabbit IgG (2729, Cell Signaling Technology, 1:5000) were also used. Target antigens were visualized using SuperSignal™ West Pico PLUS chemiluminescent substrate (Thermo Fisher Scientific, Waltham, MA, USA). Images were scanned with a ChemiDoc Imaging System (Bio-Rad, Hercules, CA, USA). Representative images are shown throughout the study.

### 2.6. Histone extraction

Histones were isolated as described [24,44]. In brief, cells were lysed using a buffer composed of 10 mM Tris–HCl (pH 7.4), 10 mM NaCl, and 3 mM MgCl_2_. After centrifugation at 2,500 rpm for 10 min at 4°C, the supernatant was discarded, and the pellets were re-lysed in the same buffer with the addition of 0.5% NP40 on ice for 10 min with gentle agitation. The nuclei were then pelleted by centrifugation at 2,500 rpm for 10 min at 4°C and resuspended in a solution of 5 mM MgCl_2_ and 0.8 M HCl. This suspension was incubated on ice for 30 min to facilitate histone extraction. Following incubation, the samples were centrifuged at 14,000 rpm for 10 min at 4°C to remove debris, and the supernatants were transferred to a clean tube. To precipitate the histones, 50% trichloroacetic acid was added. The resulting pellets were washed with acetone, air-dried, and then resuspended in a solution of 100 mM Tris–HCl (pH 7.5), 1 mM EDTA, and 1% sodium dodecyl sulfate (SDS). The concentration of histones in the extract was determined using the BCA assay (Pierce Technologies, Rockford, IL) according to the manufacturer’s instructions. Histone extracts underwent Western Blot analysis and the primary antibodies used were as follows: anti-total H3 (05–928, Millipore, 1:2000), anti-H3R17me2a (07-214, Upstate, 1:1000), anti-total-H4 (ab10158, Abcam, 1:1000), anti-H4R3me2a (39705, Active Motif, 1:1000).

### 2.7. ApoTox-Glo^TM^ Triplex assay kit

The ApoTox-Glo Triplex Assay was used according to the manufacturer’s protocol (Promega, Ref: G6321). Briefly, cells were seeded in 96-well plates and treated with the respective concentration of each compound for 72h and analyzed regarding cytotoxicity, viability, and caspase activation within a single-assay well. Each condition was represented by 6 wells. Cell viability and cytotoxicity were simultaneously assessed by measuring live- and dead-cell protease activities using fluorogenic substrates. Live-cell protease activity was detected with the cell-permeant substrate GF-AFC, while dead-cell protease activity was measured using the cell-impermeant substrate bis-AAF-R110. Fluorescence was measured after 1h incubation at room temperature at 400/505 nm (viability) and 485/520 nm (cytotoxicity).

Apoptosis was evaluated using the Caspase-Glo 3/7 assay. Cells were incubated with the Caspase-Glo 3/7 reagent for 30 min, and luminescence proportional to caspase-3/7 activity was recorded after background subtraction. Vehicle-treated cells were used as controls and data were normalized according to these values.

### 2.8. Comet assay and cytosolic DNA detection

Double-strand DNA (dsDNA) breaks were detected using the neutral comet assay as described previously [32]. Cytosolic DNA accumulation in cultured cells was assessed by immunoflorescence using dsDNA antibody from Abcam (ab27156) essentially as reported before [45]. Briefly, cells were fixed with ice-cold methanol for 10 min at room temperature and washed twice with PBS. Cells were permeabilized with 0,1% Triton X-100 for 10 min at RT. After washing, coverslips were blocked with 1% BSA in PBS for 1 h at room temperature and incubated overnight at 4°C with primary antibody diluted in 1% BSA in PBS. Next day after washing with 1% BSA in PBS, cells were incubated with fluorophore-conjugated secondary antibody in 1% BSA in PBS for 1h at room temperature, washed and stained with vectashield (Vector laboratories, Burlingame, CA, USA) containing DAPI. Images were obtained using the Zeiss Axio Imager.M1 microscope (Zeiss, Oberkochen, Germany).

### 2.9. Enzyme-linked immunosorbent assay (ELISA)

Cell culture supernatants were collected from experimental and control cultures at designated time points, centrifuged at 4500 rpm for 5 min at 4°C to remove cellular debris, and stored at −80°C until analysis. The concentration of CXCL10 in cell culture supernatants was determined using the Human CXCL10/IP-10 ELISA Kit (BD Biosciences 550926, Franklin Lanes, NJ, USA) following the manufacturer’s instructions.

### 2.10. RT and qPCR

Total RNA from cell lines was extracted using the automated Maxwell system from Promega. Quantitative reverse transcription PCR (qRT-PCR) was performed as reported, and gene expression was normalized relative to that of the housekeeping gene *H3F3A* as described [43]. Primers sequences are available upon request.

### 2.11. Transcriptomic analyses

Total RNA from HuCCT-1 cells treated with GSK715 6µM was extracted using the automated Maxwell system from Promega. RNA was subjected to quantity and quality control using Qubit HS RNA Assay Kit (Thermo Fisher Scientific, Waltham, MA, USA) and 4200 Tapestation with High Sensitivity RNA ScreenTape (Agilent Technologies). All RNA samples were high-quality, with RIN values higher than 8. Library preparation was performed using the Illumina Stranded mRNA Prep Ligation kit (Illumina) following the manufacturer’s protocol. All sequencing libraries were constructed from 100 ng of total RNA according to the manufacturer’s instruction. Briefly, poly(A) containing RNA molecules were selected using magnetic beads coated with poly(T) oligos. Poly(A)-RNAs were fragmented and reverse transcribed into the first cDNA strand using random primers. The second cDNA strand was synthesized in the presence of dUTP to ensure strand specificity. Resulting cDNA fragments were purified with AMPure XP beads (Beckman Coulter), adenylated at 3ʹ ends and then ligated with uniquely indexed sequencing adapters. Ligated fragments were purified, and PCR-amplified to obtain the final libraries. The quality and quantity of the libraries was verified using Qubit dsDNA HS Assay Kit (Thermo Fisher Scientific) and 4200 Tapestation with High Sensitivity D1000 ScreenTape (Agilent Technologies). Libraries were then sequenced using a NextSeq2000 sequencer (Illumina).

Transcriptomic data analyses were performed essentially as described [46,47]. The first step in the workflow involves quality control and preprocessing of the raw RNA-seq data. Adapter sequences and low-quality reads are removed using TrimGalore version 0.6.0 with Cutadapt version 1.18. Subsequently, clean reads are aligned to the reference genome using a splice-aware aligner STAR version 020201 over genome version hg38. Aligned reads are then quantified at the gene level using HTseq version 0.11.0. EdgeR version 3.28.1 for R software version 3.6.3 which requires raw read counts as input and performs normalization using the trimmed mean of M-values (TMM) method. TMM normalization accounts for library size differences and composition biases, ensuring accurate comparisons between samples.

For RNA splicing analyses, first RNA-seq quality control was performed using fastqc for each FASTQ file and multiqc to integrate the results of all the samples (http://www.bioinformatics.babraham.ac.uk/projects/fastqc/). Reads were aligned to the reference genome using STAR and gene counts were estimated using featureCounts. R was used to perform the differential expression analyses. Count values were imported and processed using edgeR [48]. Expression values were normalized using the trimmed mean of M values (TMM) method and lowly-expressed genes were filtered out. For differential splicing analyses, transcript abundance was quantified using Kallisto [49].

SUPPA2 [50] was used for the generation of the alternative splicing events from the input gene annotation file and the calculation of the magnitude of splicing change (ΔPSI) and its significance across biological conditions. The splicing events analysed included: Skipping Exon (SE), Alternative 5’/3’ Splice Sites (A5/A3), Mutually Exclusive Exons (MX), Retained Intron (RI), and Alternative First/Last Exons (AF/AL).

Gene Ontology (GO) biological process enrichment analysis [51] was performed using Gene Set Enrichment Analysis (GSEA) [52] implemented in the R package clusterProfiler (version 4.10.1) [53]. The analysis was conducted using genes identified as differentially expressed (p < 0.05), without applying any threshold for effect size. Transcriptomic data are going to be deposited in NCBI’s Gene Expression Omnibus (doi:10.1093/NAR/30.1.207) and will be accessible through GEO Series accession number.

### 2.12. Organoids culture and treatments

The isolation and culture of human healthy liver-derived cholangiocyte organoids and CCA-derived organoids (tumoroids) used in this study have been described previously [54]. Organoids were seeded in 48 multi-well plates and allowed to grow for 1-2 days. Then, GSK3368715 or the same volume of vehicle (DMSO) were added to the culture medium. Medium was changed every 3-4 days for a total of 12 days. Representative images of organoid cultures were taken at the onset and completion of treatment using a Leica stereoscope and camera with a 1.6X magnification. Organoid growth was quantitated using the ImageJ software [55].

### 2.13. Statistical analyses

All experiments were conducted using a minimum of three independent biological replicates. Results are expressed as the mean ± standard error of the mean (SEM), unless stated otherwise. Statistical analyses were performed using GraphPad Prism (version 9; GraphPad Software, San Diego, CA, USA). Data normality was evaluated using the Shapiro–Wilk test. Comparisons between two groups were analyzed using a two-tailed unpaired Student’s *t*-test for normally distributed data or the Mann–Whitney *U* test for non-normally distributed data. For comparisons involving multiple groups, one-way ANOVA followed by Tukey’s post hoc test was applied to normally distributed data, whereas the Kruskal–Wallis test followed by Dunn’s post hoc test was used for non-normally distributed data. A *p* value < 0.05 was considered statistically significant.

## 3. Results

### 3.1 PRMT1 expression in human CCA

First, we examined the expression of *PRMT1* in well-characterized transcriptomic datasets from human CCA tissues. In the cohort of Sia et al., including iCCAs [56], we found a significant upregulation of *PRMT1* gene expression compared to normal bile ducts (NBD)(**Fig. 1A**). Two distinct iCCA subclasses, termed the proliferation and inflammation classes, were previously identified in this cohort; patients within the proliferation class exhibited poorly differentiated, more aggressive tumors [56], and we observed higher *PRMT1* expression in this subgroup (**Fig. 1A**). Here we also found a positive correlation between *PRMT1* expression and that of the CCA marker *KRT7*, and an inverse correlation with the tumor suppressor gene *SOX17* [2](**Fig. 1A**). Similar observations were made in another cohort including both iCCAs and eCCAs [57], where *PRMT1* expression was increased in tumor tissues compared to NBD (**Fig. 1B**). In this cohort two tumor subclasses, 1 and 2, were established with subclass 2 being associated with increased cellular proliferation and worse survival [57]. *PRMT1* expression was significantly higher in subclass 2 (**Fig. 1B**). Moreover, when tumors were subdivided according to the expression of *S100P*, a gene robustly associated with CCA aggressiveness[58], *PRMT1* levels were higher in samples with enhanced S100P expression (**Fig. 1B**). Increasing evidence indicates that CCA evolves in an immunosuppressive TME that fosters tumor growth [11,59]. A recent study stratified iCCAs into four immune subtypes according to the expression of gene signatures characteristic of different immune cell types and fibroblasts in the TME [15,60]. We applied this classification to the iCCA transcriptomic dataset from Sia et al. we found that *PRMT1* expression was significantly higher in the immune-desert subtype, representing “colder” tumors that are less likely to respond to ICIs (**Fig. 1C**).

**Figure 1.**
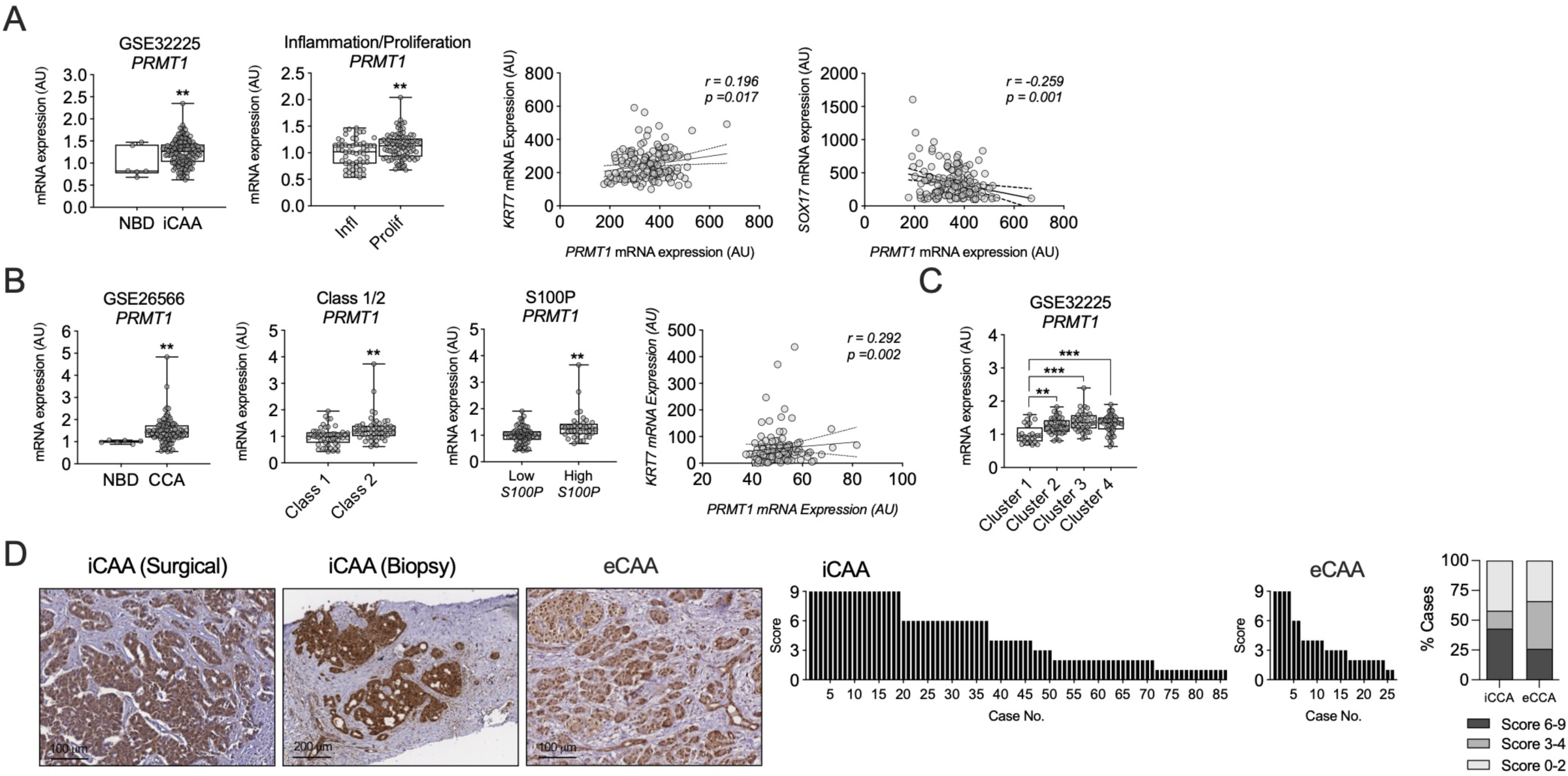
PRMT1 is overexpressed in human CCA and correlates with markers of tumor aggressiveness. **A**. *PRMT1* mRNA expression levels in intrahepatic CCA (iCCA) compared to normal bile ducts (NBD) in the GSE32225 transcriptomic dataset (left). PRMT1 expression according to the molecular subclass (Inflammation vs. Proliferation) (middle), and correlation analysis between *PRMT1* and *KRT7* or *SOX17* (right). **B**. *PRMT1* mRNA levels in CCA tissues compared to NBD in the GSE26566 dataset (left). PRMT1 levels stratified by transcriptomic class (Class 1/2) (middle) and according to the expression status of S100P, a marker of tumor aggressiveness (right). **C**. *PRMT1* expression in iCCA in GSE32225, classified according to transcriptomic TME immune subtype described by Job et al. [15]. Cluster 1: Mesenchymal; 2: Myeloid-rich; 3: immune desert; 4: reactive immunogenic stroma. **D**. Representative immunohistochemical (IHC) staining of PRMT1 in human iCCA (surgical resection and needle biopsy) and extrahepatic CCA (eCCA) tissue samples. The graphs (bottom) depict the distribution of IHC scores for each cohort. In iCCA (n=86), 43% of cases showed high expression (score 6–9), 15% medium (score 3– 4), and 42% low (score 0–2). In eCCA (n=23), 26% showed high expression, 40% medium, and 34% low expression. Data are presented as mean ± SEM. p-values are indicated in the graphs.

PRMT1 expression was also examined by immunohistochemistry, and it was readily detected in malignant tissues (both iCCA and eCCA), including iCCA needle biopsies (patients’ and tumors’ characteristics are described in **Table S1**), and quantitative evaluation encompassing signal intensity and the proportion of positive cells stained indicated that both iCCAs and eCCAs frequently overexpress PRMT1 (**Fig. 1D**) (representative images are shown in **Fig. S1**).

### 3.2 Effect of PRMT1 targeting on the growth of human CCA cells

An initial indication that PRMT1 represents a relevant therapeutic target in CCA was obtained through analysis of CRISPR/Cas9 dropout screen data from human CCA cell lines (n=28) available in the Sanger DepMap database(https://score.depmap.sanger.uk/) [61]. Genetic inactivation of *PRMT1* significantly impaired cell growth across all CCA cell lines examined (**Fig. 2A**). Next, we tested the effect of three PRMT1 (type I PRMT) pharmacological inhibitors, GSK3368715, TC-E5003 and MS023 [33,39,62], on the growth of three well-characterized CCA cell lines [40,63]. Cell growth inhibition was observed in all cases at low micromolar doses of these molecular probes (**Fig. 2B**). In agreement with this antiproliferative activity, PRMT1 inhibition also markedly impaired the clonogenic potential of CCA cells (**Fig. 2C**).

**Figure 2.**
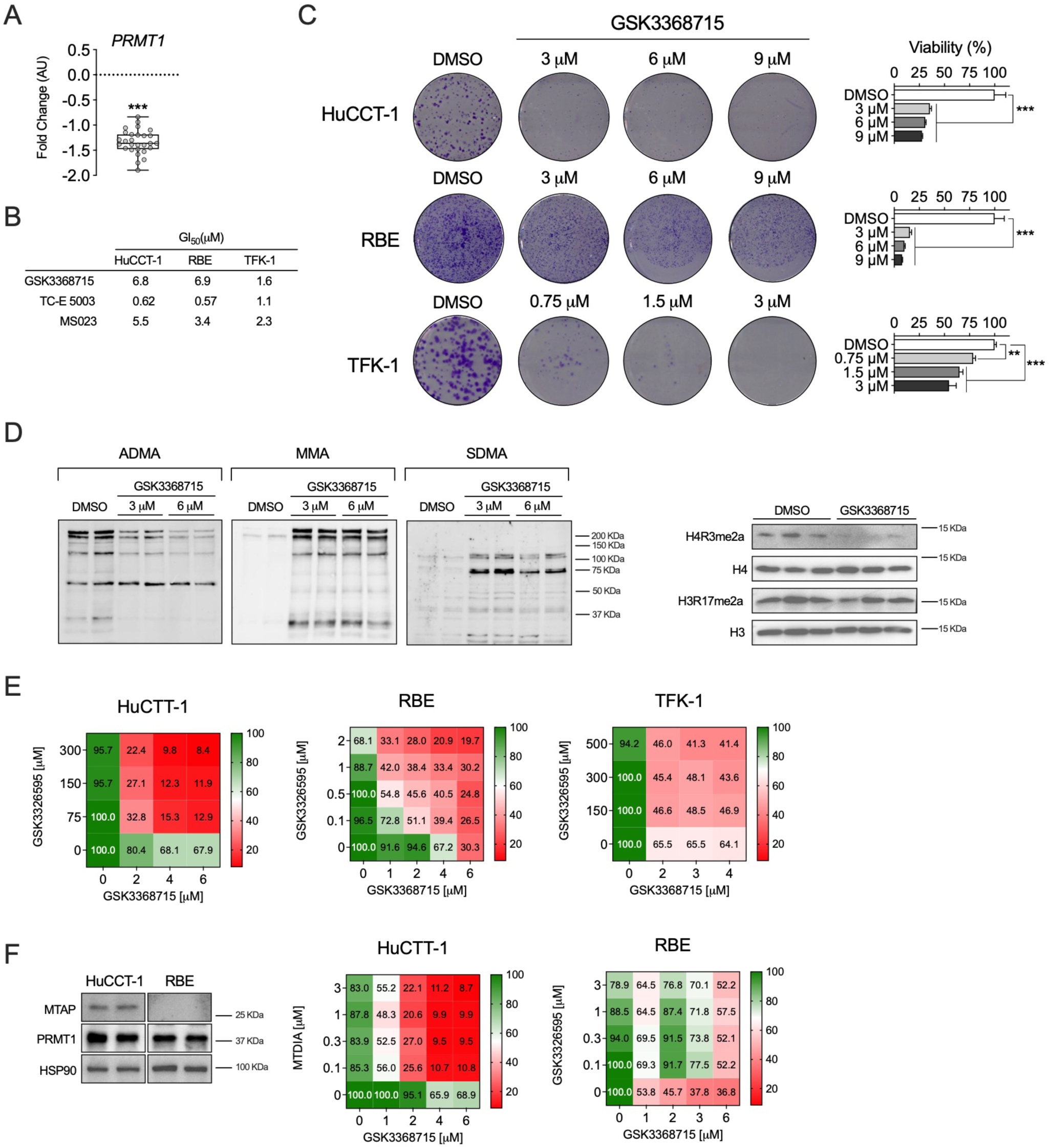
PRMT1 is essential for CCA cell survival and its pharmacological inhibition impairs proliferation and induces apoptosis. **A**. CRISPR/Cas9 gene dependency scores for PRMT1 in CCA cell lines, retrieved from the DepMap portal (Sanger Institute). Negative fitness scores indicate reduced cell viability upon *PRMT1* knockout. **B**. Table summarizing the half-maximal growth inhibitory concentration (GI_50_, μM) of three PRMT inhibitors: GSK3368715 and MS023 (Type I inhibitors), and TC-E 5003 (PRMT1-specific), across three CCA cell lines (HuCCT-1, RBE, and TFK-1) after 7 days of treatment. **C**. Representative images of colony formation assays (left) and quantification of cell viability (right) in HuCCT-1, RBE, and TFK-1 cells treated with the indicated doses of GSK3368715. **D**. Western blot analysis of global arginine methylation patterns in HuCCT-1 cells treated with GSK3368715 (0, 3, and 6 μM) for 3 days. Membranes were probed for asymmetric dimethylarginine (ADMA), monomethylarginine (MMA), and symmetric dimethylarginine (SDMA). Immunoblot validation of specific histone methylarginine marks. Levels of H4R3me2a (PRMT1-dependent) and H3R17me2a (PRMT4-dependent) were assessed in HuCCT-1 cells following GSK3368715 treatment. Total H4 and H3 served as loading controls. **E**. Analysis of drug synergism in CCA cell lines treated with GSK3368715 in combination with the PRMT5 inhibitor GSK3326595 at the indicated doses. **F**. Western blot analysis of methylthioadenosine phosphorylase (MTAP) status and PRMT1 in HuCCT-1 and RBE CCA cell lines, with HSP90 as loading control, together with analysis of drug synergism between HuCCT-1 and RBE CCA treated with MTAP inhibitor methylthio-DADMe-immucillin-A (MTDIA) and Western blot analysis of SDMA levels after MTDIA treatment. PRMT5 inhibitor JNJ64619178 was used as positive control.

To test the on-target effects of type I PRMT inhibition we first evaluated the global levels of ADMA, monomethyl arginine (MMA) and symmetric dimethyl arginine (SDMA) in HuCCT-1 and RBE cells treated with GSK3368715. We found a robust decrease in ADMA levels at all concentrations tested, and this effect was accompanied by an increase in MMA and SDMA levels (**Fig. 2D**). This response can be expected upon effective inhibition of type I PRMT activiy due to the sequential mechanism of ADMA generation, which is preceded by the monomethylation reaction, and the overlapping substrate specificity and substrate competitive behaviour between PRMT1 and PRMT5 (the main effector of SDMA) enzymes, which scavenge each other’s substrates [27,64,65]. Consistent with these observations, we found that asymmetric dimethylation of H4R3 (H4R3me2a), a key chromatin modification catalyzed by PRMT1 linked to transcriptional activation[66], was reduced upon treatment with GSK3368715, while H3R17me2a levels, a mark mainly mediated by PRMT4-CARM1 [67], were unaffected (**Fig. 2D** and **Fig. S2**).

Consistent with the functional interplay between PRMT1 and PRMT5 described above, combined inhibition of both enzymes has previously been shown to exert synergistic antitumoral effects [64,68]. In line with these findings, we also observed an enhanced antiproliferative effect in CCA cells treated with both PRMT1 and PRMT5 inhibitors, particularly in HuCCT-1 cells (**Fig. 2E**). Notably, this synergism has been reported to be more pronounced in methylthioadenosine phosphorylase (*MTAP*)-deficient tumors [64,68], which accumulate elevated levels of the MTAP substrate, 2-methylthioadenosine (MTA), a relatively selective endogenous inhibitor of PRMT5 [69]. Accordingly, *MTAP* loss is increasingly recognized as a biomarker of sensitivity to PRMT5 inhibitors [70] and may also predict responsiveness to PRMT1 inhibition [64,68,71]. However, the prevalence of *MTAP* deletion varies across solid tumors, including CCA [32,72], which may limit the overall efficacy of PRMT5 and PRMT1 inhibitors. To circumvent this limitation, highly specific inhibitors of MTAP enzymatic activity, such as MTDIA, have been developed to induce an MTAP-null-like phenotype in MTAP-expressing cancer cells. MTAP inhibition results in intracellular accumulation of MTA and consequent inhibition of PRMT5 activity [73]. Accordingly, we observed reduced SDMA leves following MTDIA treatment in MTAP positive cells (HuCCT-1), whereas no reduction was detected in RBE cells which are MTAP-null, while the PRMT5 inhibitor JNJ64619178 was effective in both cell lines (**Fig. 2F**). This approach enhances the antitumoral efficacy of PRMT5 inhibitors and potentially that of PRMT1 inhibitors [73,74]. Consistent with this rationale, we observed a strong growth-inhibitory effect when MTDIA was combined with GSK3368715 in MTAP-expressing HuCCT-1 cells, but not in RBE cells (**Fig. 2F**). These findings suggest that this strategy may also be applicable to CCAs with intact MTAP expression.

### 3.3. Mechanisms involved in the anti-CCA effect of PRMT1 inhibition

To comprehensively and unbiasedly investigate the antitumor mechanisms associated with PRMT1 targeting, we performed transcriptomic profiling of the well-characterized HuCCT-1 iCCA cell line [40,75] following treatment with GSK3368715. Consistent with the established role of PRMT1 in regulating chromatin architecture and gene expression [33,66], we identified 1,547 genes as significantly differentially expressed, 924 upregulated and 623 downregulated (Log FC< −1 or > 1, P < 0.05) (**Fig. 3A**). Among the upregulated genes were those encoding pro-inflammatory factors, including *S100A8*, *S100A9*, *CXCL10*, and *PPBP*. In contrast, genes associated with cell growth, survival, and dedifferentiation, such as *LCN2*, *ZEB1*, *SHANK2*, *PORCN*, *HDAC4*, *BCL2*, and *EGFR*, were downregulated. Notably, expression of the tumor suppressor *NDRG2* and the pro-apoptotic factor *ATG9B* was increased (**Fig. 3A**). PRMT1 is also known to play a key role in regulating the activity of splicing factors, many of which are implicated in carcinogenesis [76,77]. Accordingly, analysis of global alternative splicing (AS) changes in our RNA-seq dataset revealed that PRMT1 inhibition induces widespread alterations in AS events. Specifically, we identified 1,480 AS events in protein-coding genes, with the majority (1,173) corresponding to skipped exons (SE) (**Fig. 3B**), consistent with the role of PRMT1 in promoting exon inclusion in cassette exons [77].

**Figure 3.**
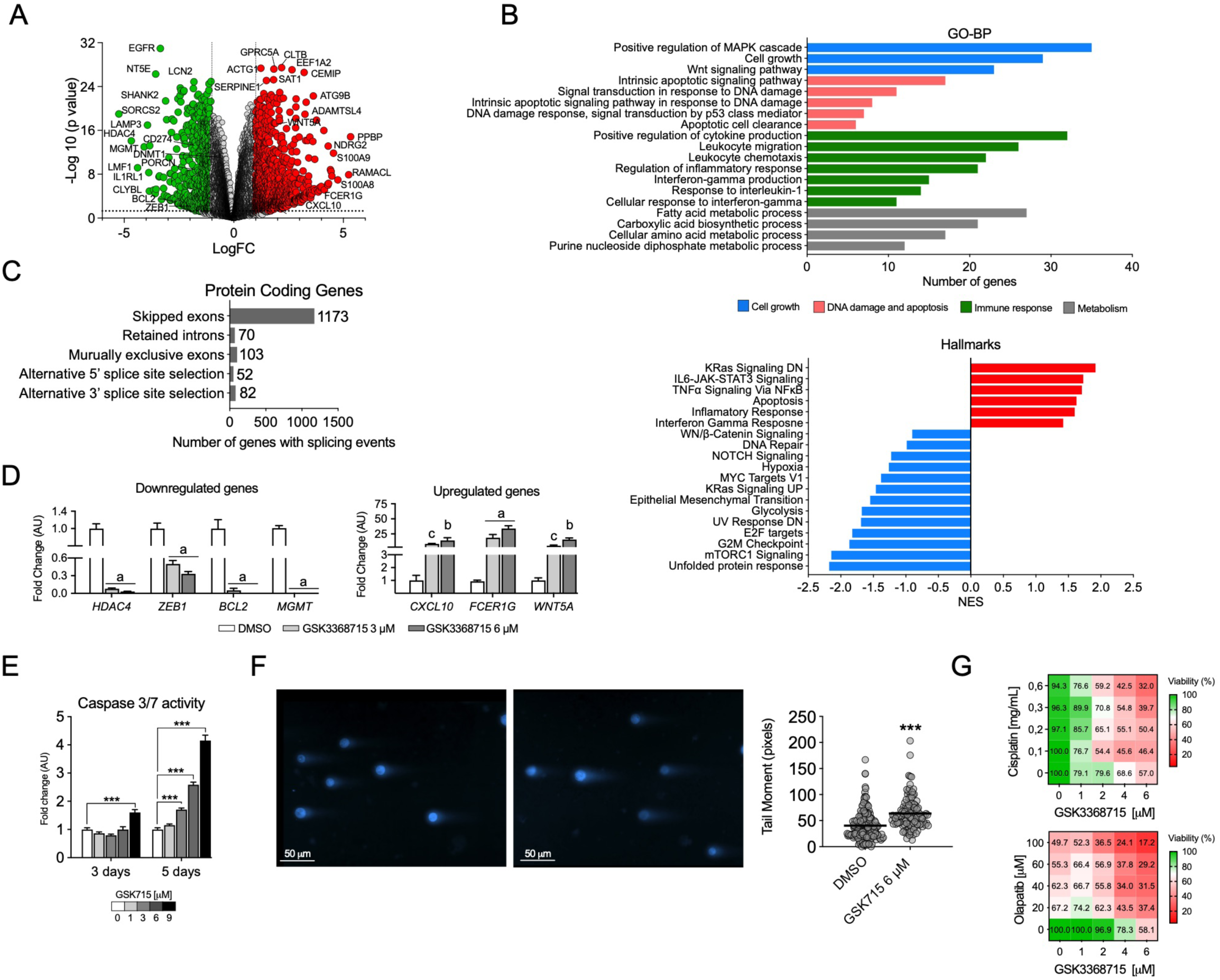
PRMT1 inhibition induces a pro-inflammatory and pro-apoptotic transcriptomic shift and sensitizes CCA cells to DNA damage response-targeting agents. A. Volcano plot displaying differentially expressed genes (DEGs) in HuCCT-1 cells treated with GSK3368715 (6 μM) for 3 days compared to vehicle control. Red dots represent significantly upregulated genes and green dots represent downregulated genes (p < 0.05). **B**. Analysis of alternative splicing events occurring after PRMT1 inhibition, including: Skipped Exons (SE), Alternative 5’/3’ Splice Sites (A5/A3), Mutually Exclusive Exons (MX), Retained Intron (RI), and Alternative First/Last Exons (AF/AL). **C**. Functional classification of DEGs using Gene Ontology Biological Processes (GO-BP). Bars represent the number of genes enriched in key functional categories: Cell growth (blue), DNA damage and apoptosis (orange), Immune response (green), and Metabolism (grey). Gene Set Enrichment Analysis (GSEA) showing the normalized enrichment scores (NES) for the top perturbed Hallmark gene sets. Red bars indicate activated pathways (positive NES) and green bars indicate suppressed pathways (negative NES) upon PRMT1 inhibition. **D**. RT-qPCR validation of selected downregulated (*HDAC4*, *ZEB1*, *BCL2*, *MGMT*) and upregulated (*CXCL10*, *FCER1G*, *WNT5A*) genes in HuCCT-1 cells treated with GSK3368715 at indicated doses. Data are normalized to *GAPDH* and presented as fold change relative to DMSO. Data are presented as mean ± SEM (n=3). a, p<0.001; b, p<0.01; c, p<0.05 vs controls. **E**. Apoptosis assessment by Caspase-3/7 activity measurement in HuCCT-1 cells treated with GSK3368715 for 3 and 5 days. Data are expressed as fold change relative to vehicle control (DMSO). Results are presented as mean ± SEM. p-values are indicated in the graphs. **F**. Representative images of comet assays showing levels of overall DNA strands breaks in HuCCT-1 cells treated with GSK3368715 for three days. Graph shows the quantification of the comet tail length (tail moment) at the level of individual cells in the number of cells indicated **G**. Analysis of drug synergism in HuCCT-1 cells treated with GSK3368715 in combination with cisplatin or olaparib at the indicated doses.

Functional classification of differentially expressed genes (DEGs) using Gene Ontology Biological Processes (GO-BP) revealed enrichment in pathways related to cell growth, DNA damage and apoptosis, immune response, and metabolism (**Fig. 3C**). Consistently, Gene Set Enrichment Analysis (GSEA) identified Hallmark pathways with positive normalized enrichment scores (NES) associated with inflammatory signaling, interferon response, and apoptosis. In contrast, pathways with negative NES values included those related to growth factor and oncogene-mediated signaling, cancer cell proliferation, hypoxia, metabolism, and DNA damage repair (**Fig. 3C**). To validate the role of PRMT1, we assessed its impact on the expression of selected genes representative of cell growth, survival, and inflammatory processes in CCA cells treated with GSK3368715 (**Fig. 3D** and **Fig. S3**). In line with the established pro-survival function of PRMT1 in cancer [33,39], inhibition with GSK3368715 led to increased caspase-3/7 activity, indicating apoptosis initiation (**Fig. 3E**). In agreement with these observations, we found that PRMT1 inhibition resulted in an increased frequency of DNA strand breaks as evaluated in a comet assay (**Fig. 3F** and **Fig. S3**). These findings suggest that targeting PRMT1 may enhance the cytotoxic effects of both standard (cisplatin) and emerging (PARP inhibitors) DNA-damage inducing therapeutic agents in CCA [78,79]. Consistent with this hypothesis, treatment of CCA cell lines with GSK3368715 in combination with either cisplatin or the PARP1 inhibitor olaparib resulted in a synergistic reduction in cell viability (**Fig. 3G** and **Fig. S3**).

To further investigate the antitumoral effects of PRMT1 inhibition, we utilized human CCA-derived organoids (tumoroids), which retain the histological features, gene expression profiles, genomic landscape, and neoplastic behavior of their tumors of origin [54]. GSK3368715 exhibited a significant growth-inhibitory effect in two tumoroid lines, CCA1T and CCA5T, with GI₅₀ values in the low micromolar range (6.4 and 3.1 μM, respectively). PRMT1 inhibition reduced tumoroid growth in a dose-dependent manner, leading to smaller organoids containing dying cells (**Fig. S4**). Notably, treatment with GSK3368715 significantly potentiated the growth-inhibitory effects of cisplatin, olaparib, and the PRMT5 inhibitor JNJ64619178 in both CCA tumoroid models (**Fig. 4**).

**Figure 4.**
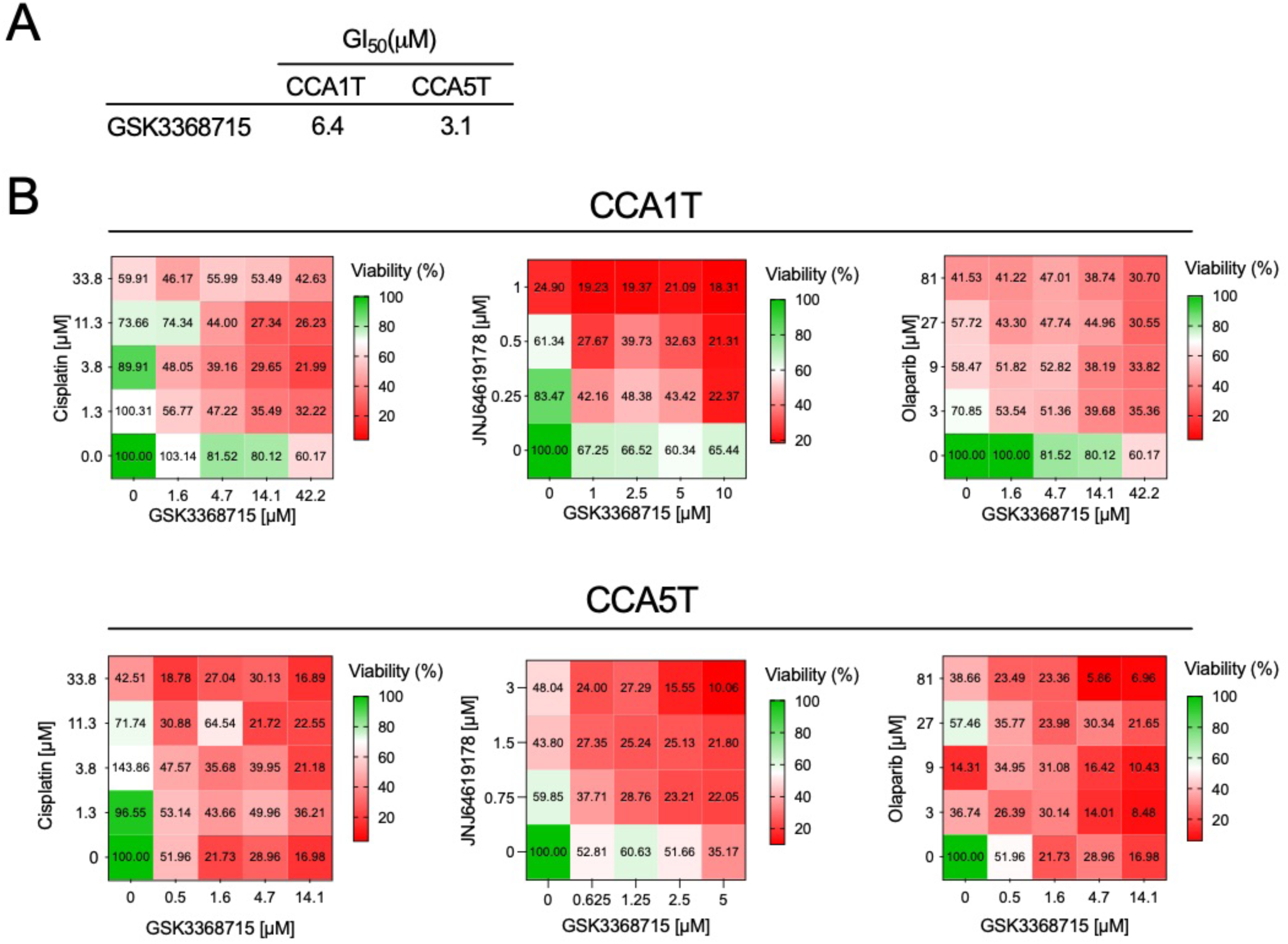
Pharmacological targeting of PRMT1 inhibits the growth of human CCA tumoroids. **A**. GI_50_ values of GSK3368715 in CCA tumoroid lines CCA1T and CCA5T. **B**. Growth inhibition of tumoroids treated with GSK3368715 in combination with cisplatin, olaparib, or the PRMT5 inhibitor JNJ64619178, showing enhanced sensitivity to combination treatments.

### 3.4. PRMT1 inhibition promotes immune-stimulated gene expression and cGAS-STING signaling in CCA cells

Gene expression analyses of CCA cells following PRMT1 inhibition revealed enrichment of pathways associated with immune responses and interferon signaling. This finding may have significant therapeutic implications, as strategies that remodel the predominantly cold and immunosuppressive tumor microenvironment in CCA could enhance the efficacy of ICI-based immunotherapy [11,14,80]. A closer look at the genes upregulated upon PRMT1 inhibition highlighted inflammatory cytokines, genes involved in DNA repair and genes associated with the interferon response (**Fig. 5A**). We validated the stimulatory effects of PRMT1 inhibition on the expression of selected genes, such as the interferon (IFN)-responsive chemokines *CXCL10* and *CXCL9*, a response that was enhanced in the presence of IFNγ (**Fig. 5B** and **Fig. S5**). Noteworthy, we also validated the upregulation of stimulator of interferon gene (STING), a key component of innate immune sensing (**Fig. 5C**). Together with cyclic GMP-AMP synthase (cGAS), STING detects cytosolic double-stranded DNA (dsDNA) and recruits and activates TANK-binding kinase 1 (TBK1), ultimately triggering phosphorylation and activation of the transcription factor interferon regulatory factor 3 (IRF3), as well as other transcription factors such as NF-κB, thereby inducing the expression of type I interferons (IFNs) and inflammatory cytokines. [81]. Consistent with the activation of this pathway, PRMT1 inhibition enhanced TBK1-IRF3 signaling elicited by IFNγ, supporting activation of the canonical cGAS-STING signaling cascade (**Fig. 5D** and **Fig. S5**). These findings were further validated when PRMT1 inhibition was combined with stimulation by dsDNA using a 70–base pair dsDNA (V70mer)[41,82] (**Fig. S5**). To further investigate the upstream events driving this response, we evaluated the accumulation of cytosolic dsDNA. PRMT1 inhibition increased cytosolic dsDNA levels (**Fig. 5E**), suggesting that genomic stress induced by PRMT1 targeting may trigger cGAS activation. Collectively, these observations indicate that PRMT1 inhibition promotes tumor-intrinsic activation of the cGAS–STING–TBK1– IRF3 signaling axis, leading to enhanced interferon signaling and induction of chemokines involved in T-cell recruitment

**Figure 5.**
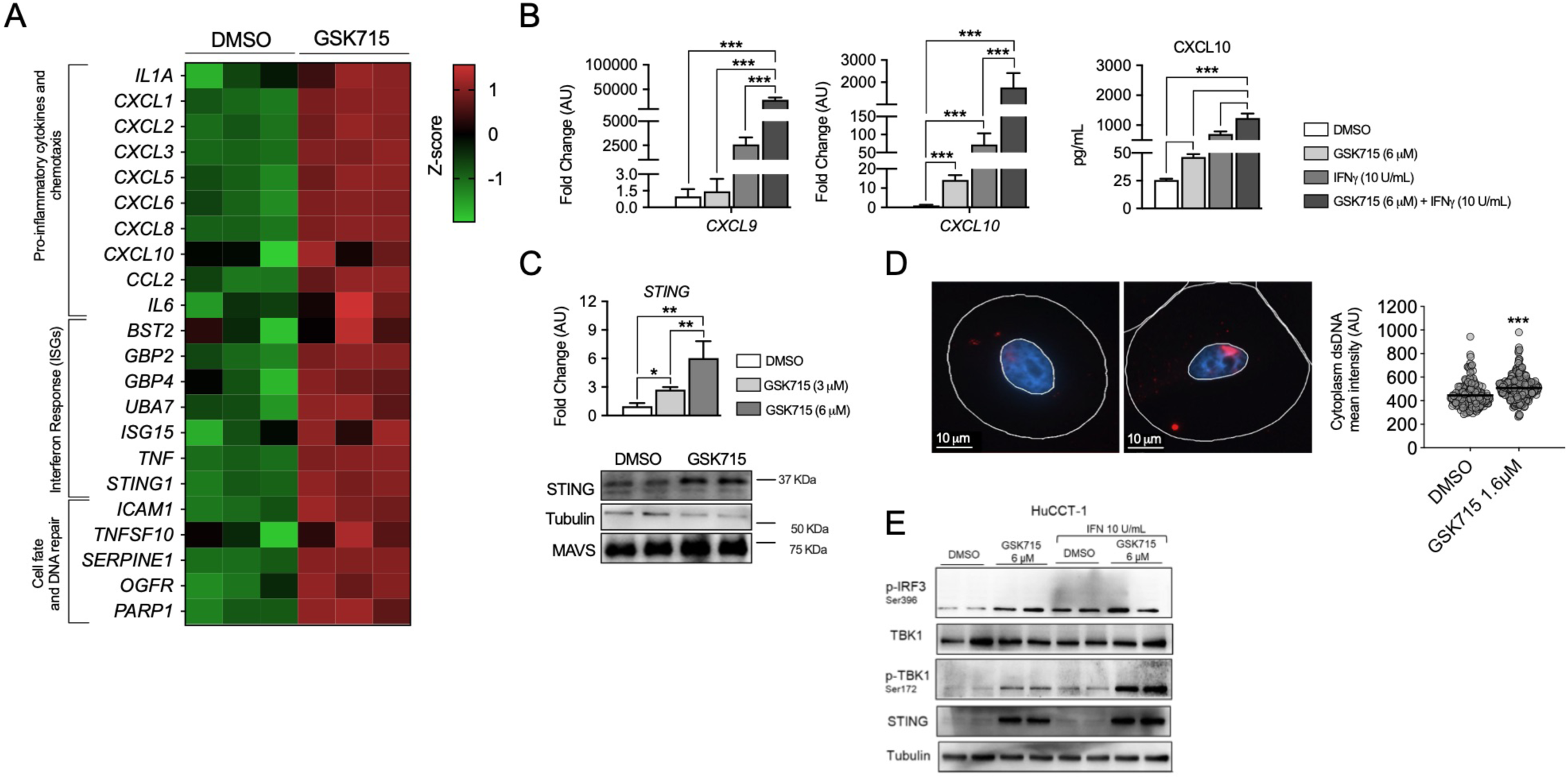
PRMT1 inhibition enhances tumor immunogenicity through activation of interferon signaling and the cGAS-STING pathway in CCA cells. A. Heatmap showing the expression of pro-inflammatory cytokines, chemokines, interferon-stimulated genes (ISGs), and genes involved in cell fate and DNA repair in HuCCT-1 cells treated with GSK3368715 compared to DMSO control. Data are represented as Z-scores. **B**. RT-qPCR analysis of selected immune-related genes (*CXCL9* and *CXCL10*) in HuCCT-1 cells treated with GSK3368715 (6 µM), IFNγ (10 U/mL), or the combination, together with ELISA quantification of CXCL10 secretion under the indicated treatment conditions. RT-qPCR data are expressed as fold change relative to control. **C**. RT-qPCR analysis of STING expression in HuCCT-1 cells treated with GSK3368715. Western blot analysis of STING protein levels in HuCCT-1 cells following PRMT1 inhibition, with Tubulin as loading control. **D**. Immunofluorescence staining of dsDNA (red) along with quantification of dsDNA signal in the cytoplasm in HuCCT-1 cells untreated or after PRMT1 inhibition with GSK3368715 (6 µM). Nuclear DNA was stained with DAPI (blue). **E**. Western blot analysis of the cGAS/STING pathway after 3 days of treatment with GSK3368715 and 24h stimulation with IFNγ (10 U/mL) in HuCCT-1 cells with Tubulin as loading control. Data are presented as mean ± SEM and p-values are indicated in the graphs. a, p<0.001; b, p<0.01; c, p<0.05 vs controls.

### 3.5. PRMT1 inhibition synergizes with PD-1 blockade to drive antitumor immunity in iCCA

We next sought to evaluate the therapeutic impact of PRMT1 inhibition in combination with ICIs in a clinically relevant TAZ/AKT-driven iCCA mouse model generated by HTVI of oncogenic plasmids [42]. This very aggressive model is based on the co-expression of constitutively active TAZ and AKT in hepatocytes, leading to the rapid and reproducible development of liver tumors that recapitulate key histopathological and molecular features of human iCCA [32,42]. Mice were assigned to receive vehicle, the PRMT1 inhibitor GSK3368715, anti-PD-1 antibodies, or their combination, and were treated for up to 3 weeks post-HTVI (**Fig. 6A**). We confirmed robust overexpression of PRMT1 in tumor tissues, supporting the rationale of targeting this pathway (**Fig. 6A**). At the end of treatments, extensive macroscopic tumor lesions were observed in control TAZ-AKT mice, whereas these lesions were markedly reduced in combination-treated animals. All treatments were well tolerated, with no significant differences in body weight observed across experimental groups (**Fig. 6B**). Strikingly, combined treatment with GSK3368715 and anti-PD-1 antibodies resulted in a significant reduction in tumor burden compared to either monotherapy or control groups, highlighting a potential synergistic antitumor effect (**Fig. 6C**). Consistently, histological analyses revealed a marked reduction in tumoral lesions area in mice receiving the combination treatment (**Fig. 6D**).

**Figure 6.**
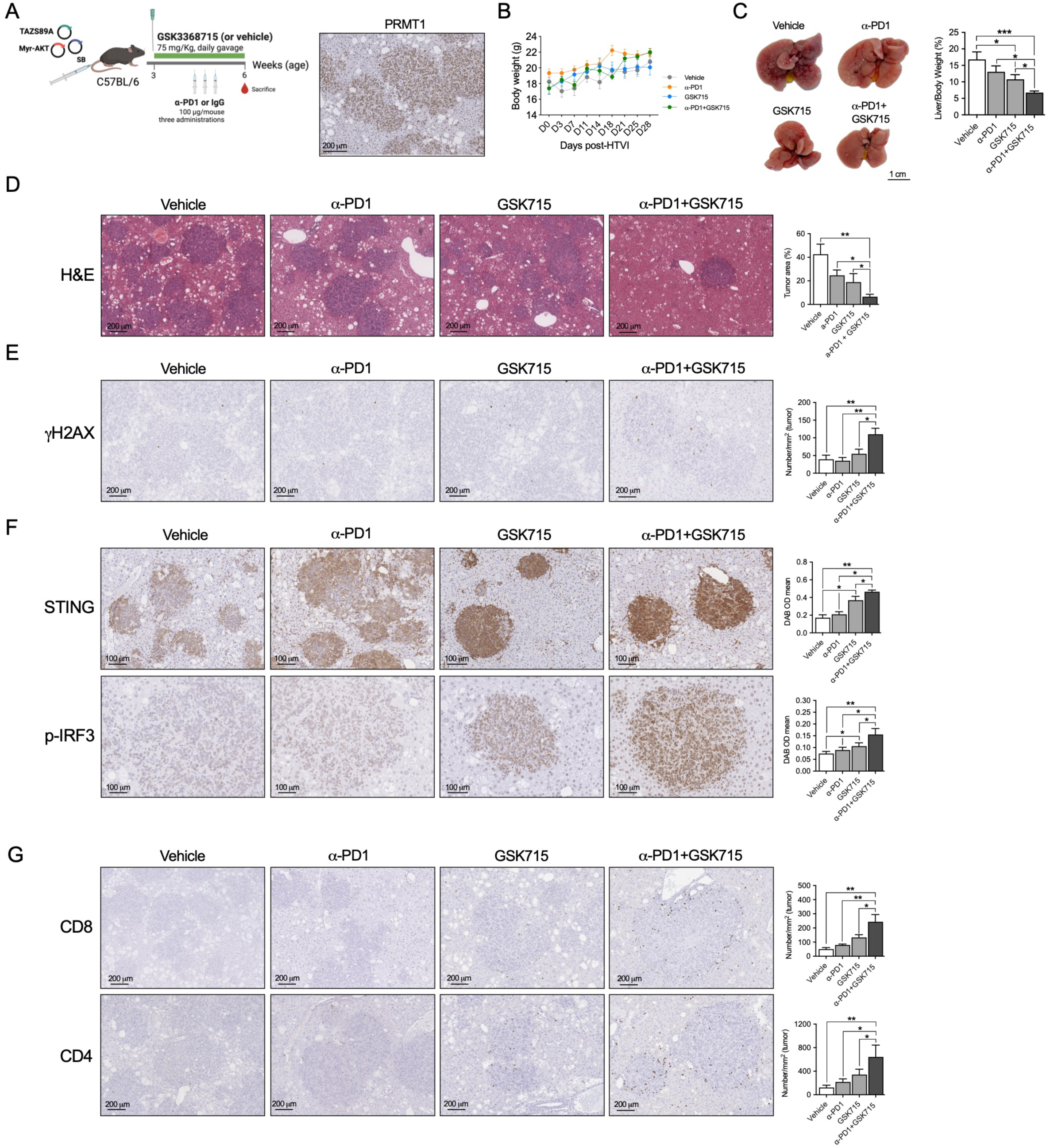
Antitumoral effects of PRMT1 inhibition in an aggressive CCA model driven by TAZ and AKT activation. **A**. Schematic representation of the hydrodynamic tail vein injection (HTVI) model based on co-expression of activated TAZ and AKT, and experimental treatment schedule (n=6 mice per group). Representative immunohistochemical (IHC) staining of PRMT1 expression in liver tumor sections from TAZ/Akt model. **B**. Body weight monitoring of mice throughout the treatment period. **C**. Tumor burden assessment shown as liver-to-body weight ratio (liver index), together with representative macroscopic images of livers from each treatment group. **D**. Representative hematoxylin and eosin (H&E) staining of liver sections showing tumor lesions, with quantification of tumor area. **E**. Detection of DNA damage by γH2AX immunohistochemistry in tumor sections from the different treatment groups, with quantification of γH2AX-positive cells. **F**. IHC detection of STING and phosphorylated IRF3 (p-IRF3), including nuclear localization of p-IRF3, consistent with activation of the cGAS-STING signaling pathway, with corresponding quantification. **G**. Representative images and quantification of tumor-infiltrating CD4⁺ and CD8⁺ T cells in control, anti-PD1, GSK3368715, and combination-treated mice. Data are presented as mean ± SEM. p-values are indicated in the graphs.

At the mechanistic level, combination-treated tumors exhibited increased γH2AX staining, consistent with elevated DNA damage (**Fig. 6E**). In agreement with our *in vitro* findings, this was accompanied by upregulation of STING and phosphorylated IRF3 (p-IRF3) in tumoral lesions (**Fig. 6F**). Notably, nuclear accumulation of p-IRF3 was significantly enhanced in the combination group, supporting activation of the STING-IRF3 axis (**Fig. 6F**). Immunohistochemical analyses revealed a marked increase in CD4⁺ and CD8⁺ T cell infiltration in tumors from mice receiving the combination therapy, indicative of enhanced immune activation within the TME (**Fig. 6G**). Collectively, these findings suggest that PRMT1 inhibition exerts antitumor activity *in vivo* and sensitizes iCCA tumors to PD-1 blockade, potentially through the induction of DNA damage and subsequent activation of innate immune signaling pathways that enhance antitumor immunity

## 4. Discussion

The identification of novel therapeutic strategies for CCA patients, particularly those capable of enhancing the efficacy of immunotherapy, remains a major clinical need given the limited response rates to current treatments and the predominantly immunosuppressive TME characteristic of this malignancy [7,11]. In this study, we identify PRMT1 as a potential pharmacological target in CCA and provide evidence supporting its role as a key regulator of tumor growth and antitumor immunity.

We first demonstrate that PRMT1 is frequently overexpressed in human CCA across independent transcriptomic datasets and patient samples. Increased PRMT1 expression was associated with molecular subtypes characterized by enhanced proliferation and poor prognosis, consistent with previous studies in other malignancies linking PRMT1 overexpression to tumor progression and aggressive phenotypes [33,35]. Notably, we found that PRMT1 expression is enriched in immune-desert tumors, which are characterized by low immune infiltration and reduced responsiveness to ICIs [15,60]. These findings indicate that PRMT1 activity might be repressing antitumor immune responses in CCA, as has been shown for other tumor types like melanoma and breast cancer [83–85].

Functional analyses revealed that both genetic and pharmacological inhibition of PRMT1 markedly impaired CCA cell growth and clonogenic capacity, supporting a critical role for PRMT1 in maintaining tumor cell fitness. These findings are in line with previous reports showing that PRMT1 regulates key oncogenic processes including transcriptional control, RNA splicing, and DNA damage responses [66,77]. Transcriptomic analyses revealed that PRMT1 inhibition induces extensive changes in gene expression and alternative splicing, affecting pathways related to proliferation, apoptosis, metabolism, DNA damage response, and immune signaling. The downregulation of oncogenic pathways together with the upregulation of pro-apoptotic and inflammatory genes highlights the central role of PRMT1 in maintaining CCA cells homeostasis, and suggest that the antitumorigenic mechanisms of PRMT1 targeting are likely diverse. Previous studies identified an important role for PRMT1 in the maintenance of DNA integrity, being involved in the homologous recombination and non-homologous end joining repair pathways [35,86]. Although the specific mechanisms by which PRMT1 contributes to DNA damage repair in CCA cells remain to be fully defined, these findings may account for the synergistic cytotoxic effects we observed when PRMT1 inhibition was combined with cisplatin or olaparib. Our results also supported the existence of a functional dependency between PRMT1 and PRMT5 in CCA cells, highlighting the therapeutic potential of dual targeting of these enzymes. The enhanced antiproliferative effects observed following combined PRMT1 and PRMT5 inhibition are consistent with previous reports describing synthetic vulnerability in MTAP-deficient contexts [64,68]. Importantly, our findings also demonstrate that pharmacological inhibition of MTAP using MTDIA can phenocopy MTAP loss in MTAP-proficient CCA cells, leading to reduced PRMT5 activity and increased sensitivity to PRMT1 inhibition. The selective efficacy of the MTDIA and GSK3368715 combination in MTAP-expressing HuCCT-1 cells, but not in MTAP-null RBE cells, underscores the importance of MTAP status as a determinant of therapeutic response. These data suggest that combining MTAP and PRMT1 inhibition may represent a promising strategy for the treatment of CCAs retaining MTAP expression. Collectively, the data further support the potential of these combinatorial approaches to achieve a favorable therapeutic index in CCA [87], although the potential toxicities associated with these combination regimens remain to be thoroughly evaluated.

We observed that PRMT1 inhibition induced genomic stress was accompanied by increased DNA damage and accumulation of cytosolic dsDNA, which in turn activates the cGAS-STING signaling axis. The cGAS-STING pathway is a central component of tumor-intrinsic innate immune sensing, linking genomic instability to the activation of type I interferon responses and antitumor immunity [88]. In this context, epigenetic regulators have emerged as key modulators of DNA sensing pathways, as their inhibition can promote the cytosolic accumulation of endogenous nucleic acids [89,90] and enhance tumor immunogenicity through cGAS-STING activation [91]. Among them, PRMTs have emerged as important modulators of immune signaling pathways. PRMT5 has been shown to suppress cGAS-STING activation through methylation of the DNA sensor IFI16, thereby attenuating interferon responses and promoting immune evasion [41]. In parallel, emerging evidence suggests that PRMT1 can negatively regulate cGAS-STING signaling through the methylation of cGAS, limiting cytosolic DNA sensing and downstream interferon responses, ultimately contributing to immune escape mechanisms in cancer [92]. Our findings extend this paradigm by demonstrating that PRMT1 inhibition induces DNA damage and cytosolic DNA accumulation, resulting in activation of the cGAS-STING axis and upregulation of interferon-stimulated genes. Moreover, we also demonstrate that PRMT1 activity repressess STING gene expression in CCA cells. Interestingly, recent studies have shown that STING expression is associated with improved survival and enhanced antitumor immune infiltration in CCA [93–95]. These observations suggest that PRMT1 acts as a negative regulator of innate immune sensing in CCA cells, and that its inhibition may restore key immunosurveillance mechanisms, such as that mediated by the cGAS-STING pathway. Importantly, the induction of chemokines such as CXCL9 and CXCL10 further supports the notion that PRMT1 targeting promotes a more inflamed tumor microenvironment, facilitating T cell recruitment and potentially enhancing responsiveness to immune checkpoint blockade.

Importantly, our *in vivo* data demonstrate that PRMT1 inhibition significantly improved the efficacy of PD-1 blockade, leading to reduced tumor burden and increased infiltration of CD4⁺ and CD8⁺ T cells. This is particularly relevant in CCA, where resistance to ICIs is largely attributed to the immunosuppressive TME [11,59]. The increased immune infiltration observed in treated tumors is consistent with the observed increase in DNA damage (as indicated by γH2AX staining) [96], an enhanced expression of STING in tumoral lesions from GSK3368715-treated mice, and the activation of the cGAS-STING axis, as indicated by nuclear P-IRF3 levels. However, additional antitumoral effects on other cellular components of the TME, as well as direct effects of PRMT1 inhibition on tumoral cells as observed *in vitro*, cannot be excluded. Interestingly, these findings align with our previous work on PRMT5 in CCA (Elurbide et al., 2024), where inhibition of this enzyme also impaired tumor growth and modulated immune responses. However, while PRMT5 targeting predominantly affected mRNA splicing and DNA repair pathways, our data suggest that PRMT1 could play a more direct role in regulating tumor-intrinsic innate immune sensing. These observations highlight the complementary roles of different PRMT family members in cancer biology. More broadly, our study contributes to the growing body of evidence indicating that epigenetic therapies can modulate tumor–immune interactions and enhance the efficacy of immunotherapy [25,26]. In tumors such as CCA, characterized by a predominantly non-inflamed microenvironment, targeting epigenetic regulators such as PRMT1 may represent a promising strategy to overcome immune resistance. Altogether, our study identifies PRMT1 as a critical regulator of tumor growth, genomic stability, and immune evasion in CCA, and provides a strong rationale for the clinical evaluation of PRMT1 inhibitors, particularly in combination with ICIs and DNA-damaging agents for the treatment of CCA.

## Acknowledgements

The authors gratefully acknowledge the excellent technical support provided by Mr. Roberto Barbero, Ms. Miriam Belzunce, and Ms. Florida Ahmed. They also sincerely thank Mr. Eduardo Avila for his generous support, as well as the patients whose samples were used in this study.

